# *De novo* assembly of 20 chickens reveals the undetectable phenomenon for thousands of core genes on sub-telomeric regions

**DOI:** 10.1101/2021.11.05.467060

**Authors:** Ming Li, Congjiao Sun, Naiyi Xu, Peipei Bian, Xiaomeng Tian, Xihong Wang, Yuzhe Wang, Xinzheng Jia, Rasmus Heller, Mingshan Wang, Fei Wang, Xuelei Dai, Rongsong Luo, Yingwei Guo, Xiangnan Wang, Peng Yang, Shunjin Zhang, Xiaochang Li, Chaoliang Wen, Fangren Lan, AMAM Zonaed Siddiki, Chatmongkon Suwannapoom, Xin Zhao, Qinghua Nie, Xiaoxiang Hu, Yu Jiang, Ning Yang

**Affiliations:** Key Laboratory of Animal Genetics, Breeding and Reproduction of Shaanxi Province, College of Animal Science and Technology, Northwest A&F University, Yangling 712100, China; National Engineering Laboratory for Animal Breeding and Key Laboratory of Animal Genetics, Breeding and Reproduction, Ministry of Agriculture and Rural Affairs, China Agricultural University, Beijing 100193, China; State Key Laboratory of Agrobiotechnology, College of Biological Sciences, China Agricultural University, Beijing 100193, China; National Research Facility for Phenotypic and Genotypic Analysis of Model Animals (Beijing), China Agricultural University, Beijing 100193, China; Department of Animal Science, Iowa State University, Ames, IA 50011, USA; School of Life Science and Engineering, Foshan University, Foshan 528225, China; Section for Computational and RNA Biology, Department of Biology, University of Copenhagen, Copenhagen N 2200, Denmark; Howard Hughes Medical Institute, University of California Santa Cruz, Santa Cruz, CA 95064, USA; Department of Ecology and Evolutionary Biology, University of California Santa Cruz, CA 95064, USA; Department of Pathology and Parasitology, Faculty of Veterinary Medicine, Chittagong Veterinary and Animal Sciences University, Chittagong-4202, Bangladesh; School of Agriculture and Natural Resources, University of Phayao, Phayao, Thailand; Department of Animal Science, McGill University, Montreal, Quebec, Canada; Department of Animal Genetics, Breeding and Reproduction, College of Animal Science, South China Agricultural University, Guangzhou, 510642, Guangdong, China

**Keywords:** Chicken, Pan-genome, Missing genes, Nonconical DNA secondary structure, Avian evolution

## Abstract

The gene numbers and evolutionary rates of birds were assumed to be much lower than that of mammals, which in sharp contrast to the huge species number and morphological diversity of birds. It is very necessary to construct a complete avian genome and analyze its evolution.We constructed a chicken pan-genome from 20 *de novo* genome assemblies with high sequencing depth, newly identified 1,335 protein-coding genes and 3,011 long noncoding RNAs. The majority of these novel genes were detected across most individuals of the examined transcriptomes but were accidentally measured in each of the DNA sequencing data regardless of Illumina or PacBio technology. Furthermore, different from previous pan-genome models, most of these novel genes were overrepresented on chromosomal sub-telomeric regions, surrounded with extremely high proportions of tandem repeats, and strongly blocked DNA sequencing. These hidden genes were proved to be shared by all chicken genomes, included many housekeeping genes, and enriched in immune pathways. Comparative genomics revealed the novel genes had three-fold elevated substitution rates than known ones, updating the evolutionary rates of birds. Our study provides a framework for constructing a better chicken genome, which will contribute towards the understanding of avian evolution and improvement of poultry breeding.

## Introduction

The ~10,770 species of birds (Gill et al. 2020) show complex and diverse morphology and behavior, however the currently available avian genomes present a reduced rate of evolution and much lower gene numbers than those of all other tetrapods. The apparent discordance remained a major evolutionary conundrum. Some studies have shown that birds tend to have fewer genes than other tetrapods due to the large segmental deletions in birds (Lovell et al. 2014; Zhang et al. 2014), while other researchers suggested that these missing genes may not have been sequenced (Bornelov et al. 2017; Botero-Castro et al. 2017; Yin et al. 2019; Zhu et al. 2021). Using the advanced sequencing technique and methodology, the Vertebrate Genomes Project (VGP) found lots of genes were missing in previous genomes, even the previous and VGP assemblies were from the same individual animals (Kim et al. 2021; Rhie et al. 2021). It remains unclear how many genes within individual bird species and why some genes are missing in the currently available genomes.

Comprehensive analyses indicate multiple high-quality de novo genome assemblies possess more power to capture the complete set of genes, which leads to the appearance and prevalence of “pan-genome” in various species (Wong et al. 2018; Duan et al. 2019; Tian et al. 2020; Wong et al. 2020). The pan-genome of mammals are typically of the “closed” pattern with a limited number of variable genes (Duan et al. 2019; Li et al. 2019; Tian et al. 2020), which means the number of genes in mammalian species is relatively conserved. While bacteria, fungi, and plants exhibit the characteristic of “open” pattern, with the proportion of core genes size is less than 80% in many species (Golicz et al. 2019). Recent research using population resequencing data found that the core genome of chickens is only 76% (Wang et al. 2021), which puzzles us because it seems to be inconsistent with the status of chickens in evolution. As the most abundant class of tetrapod vertebrates, the *de novo* based pan-genome of avian has not yet been established, which is essential to solve many biological issues.

Chicken (*Gallus gallus*) as one of the most important farm animals plays a major role in human food production and has been widely used as a model organism in studies of developmental biology, virology, oncogenesis, and immunology (Cooper et al. 1966; Stehelin et al. 1976; Brown et al. 2003; Vogt 2011). In this study, we utilized 20 new high-quality assemblies of diverse chicken breeds to generate the first *de novo* based chicken pan-genome. A total of 159 Mb novel sequences containing 1,335 coding genes that completely absent from the chicken genome were identified, verified, and localized. Importantly, most of the novel genes actually exist in all of the chicken genomes but were prone to be missing in the DNA sequencing leaded by high proportions of tandem repeats and secondary structures. Hence, unwinding complex DNA structures should be one of the most important advances to improve the sequencing quality for the assembly of the complete avian genomes. Our study revealed that the numbers of chicken genes are comparable to those of other tetrapod vertebrates and a new pan-genome pattern of birds.

## Results

### Identification and validation of non-redundant novel sequences

Twenty chickens from four continents representing widespread indigenous chicken breeds, commercial broilers and layer lines were sampled for *de novo* genome assembly (**Fig. 1a**, **Supplementary Table 1**). Ten assemblies were constructed by integrating both PacBio (53 to 95×) and Illumina data (45 to 70×), resulting in a contig N50 size ranging from 5.89 to 16.72 Mb (**Supplementary Table 2**). Six of them were further clustered at the chromosome level by using Hi-C (112 to 125×) data (see **Methods**, **Supplementary Fig. 1–4**, **Supplementary Tables 2** and **3**). The remaining ten samples were assembled based on Illumina reads from a combination of libraries with multiple insert sizes, ranging from 500 bp to 5 Kb (with a depth of ~134 × per genome, **Supplementary Table 2**). These ten samples showed a contig N50 size ranging from 80.30 Kb to 137.59 Kb (**Supplementary Table 2**), representing high-quality Illumina genomes (Schatz et al. 2010). The completeness of the 20 assemblies was evaluated through the Benchmarking Universal Single-Copy Orthologs (BUSCO) analysis. Most (from 92.4% to 95.3%) of the 4,915 core genes in the Aves dataset were identified in the 20 assemblies, which is comparable to the percentage in the reference chicken genome (GRCg6a: 95.4%) and thus supports a high-quality genome assembly (**Fig. 1b**, **Supplementary Fig. 5**, and **Supplementary Table 4**).

**Figure 1.**
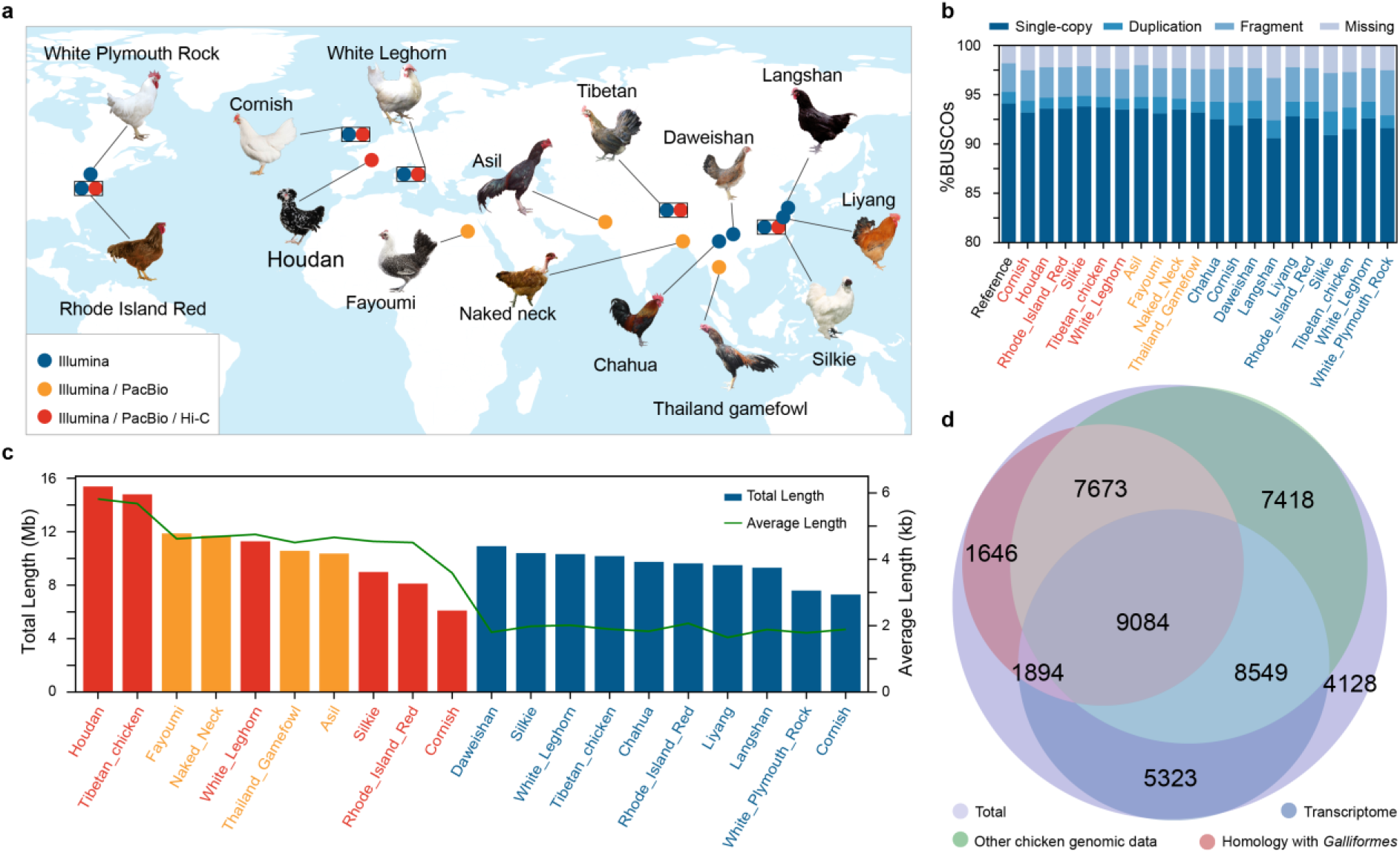
Chicken novel non-redundant sequences identified by twenty *de novo* assemblies. **a**, Geographic locations of the original chicken breeds used for *de novo* assembly and their sequencing platforms. The rectangle indicates this breed has two individuals. **b**, Genome assembly completeness assessed by BUSCO. **c**, Length of novel sequences initially obtained from 20 *de novo* assemblies. The polygonal line represents the average length and the column represents the total length. **d**, The number of novel sequences validated by other chicken genomes, homology with Galliformes and transcriptome. The colors of the breed name in **b** and **c** are consistent with **a**.

To identify novel sequences, all 20 *de novo* assemblies were aligned against GRCg6a (see **Methods**, **Supplementary Fig. 6**). For stability, we used GRCg6a from a same red junglefowl as the reference genome in the past two decades, not the newly unpublished GRCg7b from a broiler. The genome length of GRCg6a (1.06 Gb) and GRCg7b (1.05 Gb) are almost the same. Unaligned sequences or sequences with < 90% identity and > 500 bp in length to GRCg6a were retained and potentially contaminating non-Chordata sequences were removed. After these screening, each assembly left 6.10 to 15.40 Mb of novel sequences (**Fig. 1c**). We merged the novel sequences from all 20 assemblies and built a pan-genome of chicken. The pan-genome contains 158.98 Mb of non-redundant novel sequence, which obtained from 45,715 sequences with an average length of 3,478 bp. (**Table 1**, **Supplementary Fig. 7**, and **Supplementary Table 5**). The chicken pan-genome expanded the size of GRCg6a by 14.92% which is the highest percentage among the published vertebrate pan-genomes.

**Table 1.**
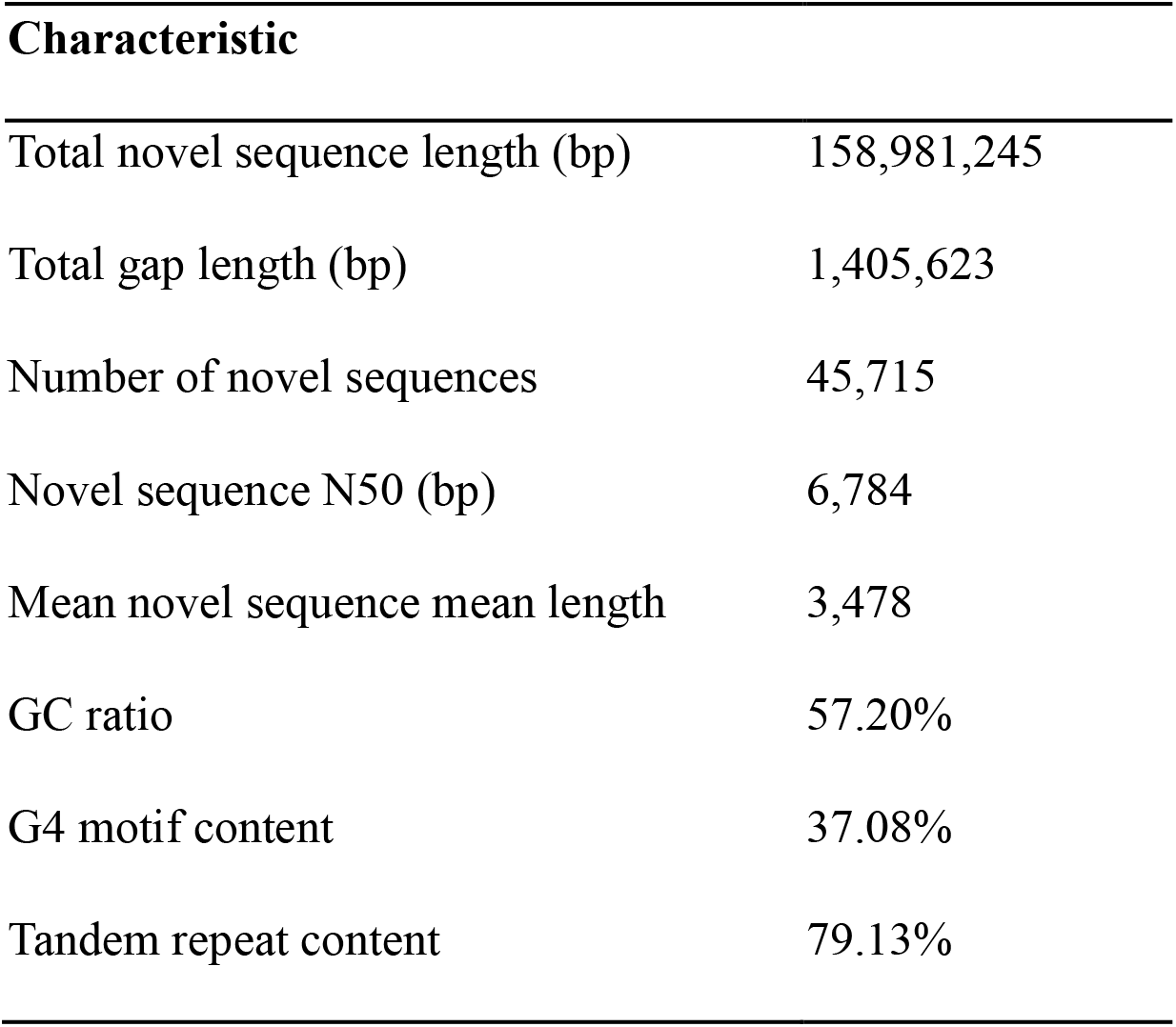
The characteristics of novel sequences in this study.

| **Characteristic** |  |
| --- | --- |
| Total novel sequence length (bp) | 158,981,245 |
| Total gap length (bp) | 1,405,623 |
| Number of novel sequences | 45,715 |
| Novel sequence N50 (bp) | 6,784 |
| Mean novel sequence mean length | 3,478 |
| GC ratio | 57.20% |
| G4 motif content | 37.08% |
| Tandem repeat content | 79.13% |

We next validate the reproduced the 158.98 novel sequences. 71.58% novel sequence can be detected in the genomes other individual, including our assemblies and 922 resequenced chicken genomes from previous studies. 44.40% can find orthologs from fourteen other publicly available *Galliformes* genomes. 54.36% can be detected from transcriptomes 263 transcriptomes from multiple tissues from 54 chickens, including 46 transcriptomes obtained from 11 tissues/organs from six individuals in our study and 217 publicly available chicken RNA sequencing (RNA-Seq) datasets from 48 individuals (Cardoso-Moreira et al. 2019) (see **Supplementary Note**, **Fig. 1d**, **Supplementary Fig. 8–10**, **Supplementary Table 5–8**, and **Supplementary Dataset**). In total, 90.97% of the novel sequences were verified in at least one of the above data sources.

### Distribution of cryptic novel sequences across chicken individuals

We found that the distribution of the novel sequences is obviously inconsistent in different verified sources. The detection rate of novel sequences in one genome is extremely low, the median is only 0.43% among the 922 resequencing data and 5% among the 20 assemblies (**Fig. 2a**). Among all 159Mb novel sequences, the 10 Illumina assemblies independently detected about 60Mb, containing only 3.44 Mb intersection with PacBio assemblies. Due to the higher detection rate of RNA-Seq, we picked up the transcribed novel sequences according to the 263 transcriptomes for further validation. RNA-Seq confirmed that 60.51% of the transcribed regions of the cryptic novel sequences were shared among more than half of the chicken genomes (**Supplementary Fig. 11**). In the six individuals with both PacBio genome assembly and transcriptome data, the transcriptomes supported a total of 16,169 novel sequences, 9,200 (56.90%) of which were detected in the transcriptomes of all six individuals. However, 5,711 (62.08%) of the 9,200 sequences were absent in all six PacBio genome assemblies (**Fig. 2b**). By mapping the PacBio reads to the novel sequences, 76.35% and 52.81% of the novel sequences were covered by at least one read across more than half or all PacBio-sequenced individuals, respectively. Moreover, although the GRCg6a assembly did not contain our novel sequences, 6.30% (2,879) of the sequences were covered by the Illumina sequencing reads of the GRCg6a individual with at least 7× coverage (corresponding to 25% of the genome-wide depth) (**Supplementary Fig. 12** and **Supplementary Dataset**). To explain the prevalence of ubiquitously transcribed yet missing novel sequences in the assemblies, we compared the median sequencing depth of the novel sequences with the whole-genome depth in the individuals. We found that the median sequencing depth of the novel sequences was only one-third of the whole-genome depth in the individuals in which the novel sequences were successfully assembled. Furthermore, in the individuals in which a given novel sequence was missing from the assembly, the median sequencing depth of the novel sequences was only one-twentieth of the whole-genome depth, which is insufficient for successful assembly (**Fig. 2c**). Collectively, the results indicated that the novel sequences were most likely present in most or all the chicken genomes but were prone to be missing in the assemblies due to their extremely low DNA sequencing depth.

**Figure 2.**
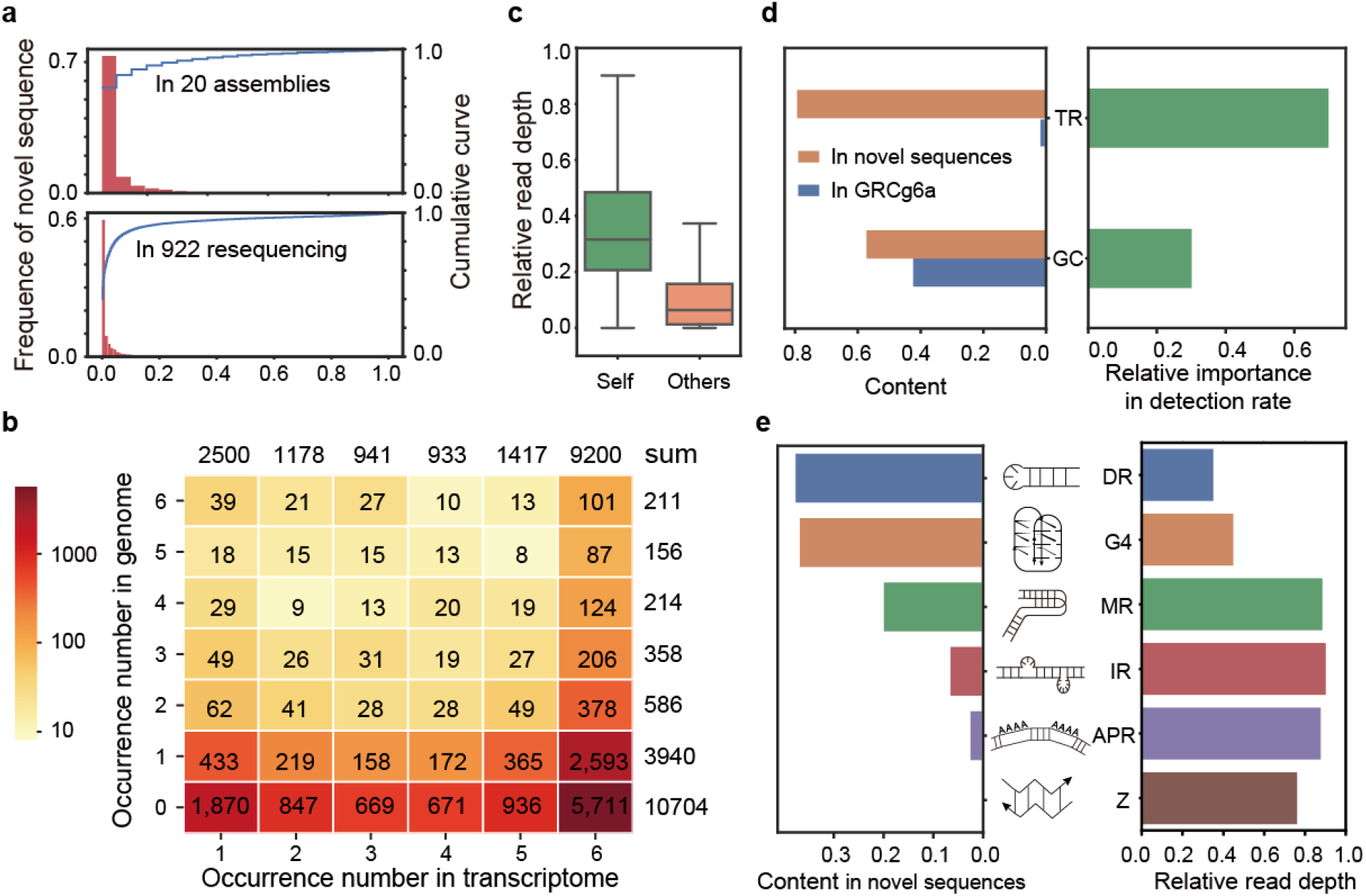
Characterization of novel sequences. **a**, The distribution and cumulative curve of observed frequencies of novel sequences in 20 assemblies and 922 resequenced individuals. **b**, The observed frequency of the expressed novel sequences in the transcriptomes of six chickens(column) and their corresponding genomes (row). **c**, Relative read depth of novel sequences in the specific assembly which the novel sequence was present (green) and absent (orange) The whole genome read depth was set to one. **d**, Left: the TR and GC content of the GRCg6a and novel sequences, respectively; Right: the feature importance of TR and GC for the detection rate of novel sequences. **e**, Left: the content of noncanonical DNA structures in the novel sequences; Middle: the putative structures of noncanonical DNA; Right: the read depth ratio of novel sequences with or without noncanonical DNA structures.

### Cryptic novel sequences have a high content of tandem repeats

We observed a higher GC content in the novel sequences than in the reference genome (57.2% vs. 42.30%). However, the GC content around 60% could not significantly reduce the depth of sequencing. Another influence of sequencing is repeat. The content of tandem repeats (TRs) in the novel sequences was 79.13%, which is extremely high and significantly higher than in GRCg6a (2.2%; chi-squared test, *P*-value = 0) (**Fig. 2d** and **Table 1**). Other interspersed repeats such as LTR and LINE were low (0.09% in novel sequence vs 9.6% in GRCg6a, **Supplementary Fig. 13**). We predicted the relative importance of TR and GC content in detection rate in assembly using random forest classifier and found the TR content had a greater influence than GC (**Fig. 2d**, **Supplementary Fig. 14**). The TR can form noncanonical DNA structures, such as G-quadruplexes (four-stranded noncanonical DNA/RNA topologies, hereafter referred to as G4 motifs), Z-DNA, A-phased repeats and inverted repeats, which can form cruciforms, triplexes and slipped structures, leading to genomic instability (Zhao et al. 2010) and incapable DNA sequencing (Guiblet et al. 2018). We found these noncanonical structure are highly intersected with TR regions (**Supplementary Fig. 15**). Among these structures, the content of direct repeats (DR) (37.96%) and G4 motifs (37.08%), are the highest in novel sequences, while their occurrence in GRCg6a is only 1.47% and 0.77%. DR and G4 also showed the largest negative correlation with read depth, the novel sequence with DR and G4 motif had only 1/3 and 1/2 read depth of all novel sequences. (**Fig. 2e**). It is worth noting that as particularly stable noncanonical DNA structures, G4 motifs typically form in guanine-rich regions of genomes, which may be one of the reasons why GC-rich sequences are difficult to sequence. We also found that the transcribed regions of novel sequences showed a lower TR content (**Supplementary Fig. 16**), which might be the reason why RNA-Seq resulted in a higher observed frequency than DNA sequencing.

### Abundant genes are embedded in novel sequences

Within the novel sequences, the expressible regions are of the most interest for potentially discovering novel candidate genes. To identify novel chicken genes, we performed gene annotation for all 20 assemblies by *de novo* and reference-guided methods using the multi-tissue transcriptomes (see **Methods**, **Supplementary Table 8**). The median expression level of these putative novel genes was significantly higher than the median expression of GRCg6a-annotated genes (*P*-value = 2.84 × 10^-7^) (**Fig. 3a** and **Supplementary Fig. 17**). The chromosome conformation analysis showed that the regions containing novel genes were significantly enriched in the A-compartment (*P*-value = 2.2× 10^-16^) (**Fig. 3b** and **Supplementary Fig. 18**), which is associated with open, expression-active chromatin. Furthermore, the orthologues of the novel genes showed expression levels that were higher than the median levels observed in other species, such as human and mouse (**Supplementary Fig. 19**), suggesting plausible functions and active expression of these genes.

**Figure 3.**
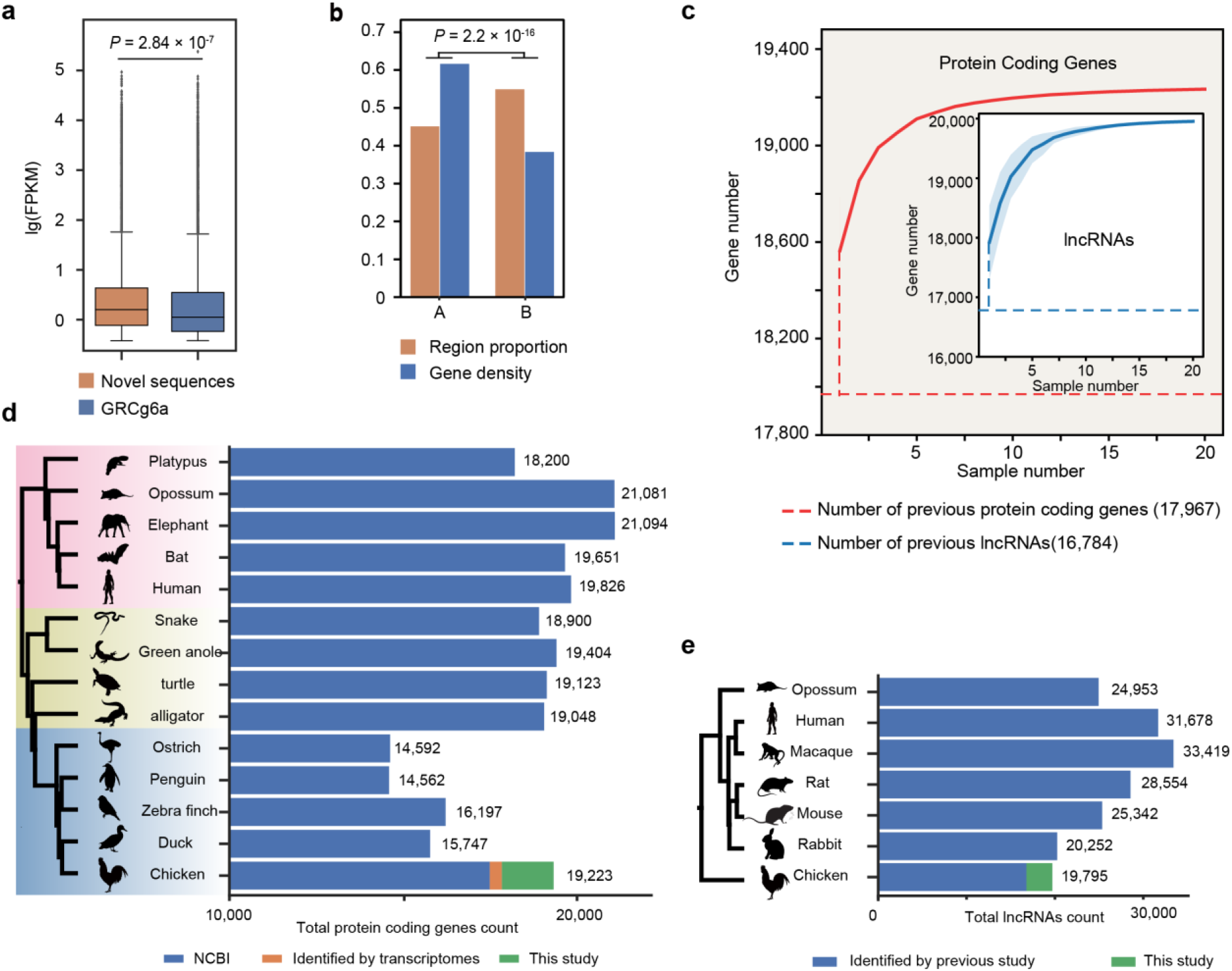
Abundant genes embedded in novel sequences. **a**, Relative expression of transcripts of reference and novel sequences, respectively. **b**, The region proportion of A/B compartment and the novel gene number proportion in A/B compartment, respectively. **c**, The identification of protein coding genes/lncRNAs increased with sample numbers. The shaded area indicates the 95% confidence interval. **d**, The number of protein coding genes in representative species including mammals, reptiles and birds. Blue, orange and green columns refer to protein coding genes identified by NCBI, Yin et al (2019). and our study, respectively. **e**, Total lncRNAs numbers of mammalian representative species and chicken. Blue and green columns refer to lncRNAs identified by Sarropoulos et al (2019). and this study, respectively.

We identified 1,335 novel coding genes with FPKM > 1, and completely missing from GRCg6a (see **Methods**, **Supplementary Fig. 20**, and **Supplementary Table 9**). The novel coding genes were distributed across 1,100 novel sequences, with an average length of 1,047 bp. By searching against the non-redundant (nr) protein database of NCBI (E-value ≤ 1×10^-5^), 969 of the novel coding genes were found to show Chordata protein orthologues, 738 of which belonged to Aves (**Supplementary Table 9**). In addition to novel coding genes, we also identified 3,874 confident transcripts which complemented 1,336 partially missing coding genes in GRCg6a (**Supplementary Fig. 21** and **Supplementary Table 10)**.

To validate the novel coding genes, proteomic analysis of multiple tissues (hypothalamus, spleen and cecal tonsil) was performed via an LC-MS/MS strategy (**Supplementary Table 8**). A total of 255 (19.10%) novel coding genes were confirmed by the existence of corresponding proteins (**Supplementary Table 9**), compared to 6,201 (35.48%) of all the coding genes present in the reference genome. The lower detection rate of novel genes in proteomics may be affected by the differences in protein length and the quality of the protein database used for searching. Notably, after removing novel coding genes less than 1 Kb in length, the proteomic verification ratio of the remaining novel coding genes increased to 29.11%.

We found that most of the novel coding genes were present and expressed in most chicken breeds. According to the DNA data, 92.47% of the novel sequences containing novel coding genes were supported by at least one PacBio read in each sample (**Supplementary Fig. 22**). According to the comparison of multi-tissue transcriptomes of six individuals, 55.13% and 80.97% of the novel coding genes were detected in all six or at least three individuals, respectively (**Supplementary Table 9**). Based on our sequencing platform, assembly strategy and annotation pipeline, the modelling of the saturation curve by iteratively randomly sampling individuals suggested that the number of novel genes detected by genome assembly did not significantly increase beyond a sample size of ten (**Fig. 3c**). A previous study (Yin et al. 2019) based on the *de novo* assembly of massive chicken transcriptomes increased the number of known chicken coding genes from 17,477 to 17,967 (**Fig. 3d**, **Supplementary Fig. 23**). According to our chicken pan-genome, we found that the total number of chicken coding genes reached at least 19,223 (**Fig. 3d**, **Supplementary Fig. 23**, and **Supplementary Table 11**).

In addition to coding genes, we identified 3,011 long noncoding RNAs (lncRNAs) (see **Methods**, **Supplementary Table 12**). Among these novel lncRNAs, 87.85% were supported by at least one PacBio read in each sample (**Supplementary Fig. 22**). In our multi-tissue transcriptomes of six individuals, 47.72% and 75.09% of novel lncRNA genes were detected in all six or at least three individuals, respectively (**Supplementary Table 12**). The increasing saturation curve of the observed novel lncRNA genes was similar to that of novel coding genes (**Fig. 3c**). Using the same pipeline as in a previous study (Sarropoulos et al. 2019), we showed that the total number of chicken lncRNAs was at least 19,795 (**Fig. 3e**). Therefore, our study revealed that the numbers of both the protein-coding and lncRNA genes of chicken are comparable to those of other tetrapod vertebrates (**Fig. 3d** and **3e**).

### Novel sequences and genes are concentrated in sub-telomeric regions with elevated substitution rates

We anchored the novel sequences to GRCg6a based on flanking sequence alignment and chromosome interaction mapping (see **Methods**). A total of 27,966 (61.17%) novel sequences containing 1,043 novel coding genes and 1,567 novel lncRNAs were anchored to GRCg6a by at least one end (**Supplementary Table 5, 9 and 12**). Among these sequences, 6,735 novel sequences containing 388 coding genes were fully anchored by both ends. The fully anchored novel sequences were further classified as insertions, alternate alleles, or multiple alternative alleles (**Supplementary Fig. 24b**, **c**, and **d**) and were dispersed on every chromosome of GRCg6a, filling 72 of 946 gaps in GRCg6a (**Supplementary Fig. 24a**, **e**).

The fully anchored novel sequences and genes were overrepresented on micro-chromosomes (GGA11-38) (< 10 Mb) or the terminal 5 Mb ends of macro-chromosomes (**Fig. 4a**, **Supplementary Fig. 25**), which are termed as sub-telomeric regions. By comparison with the random distribution, we estimated 2.5- (*P*-value < 1 × 10^-6^, permutation) and 5-fold (*P*-value < 1 × 10^-6^, permutation) increases in fully anchored novel sequences and gene density within sub-telomeric regions, respectively (**Supplementary Fig. 26**). The novel sequences nearly doubled the length of the micro-chromosomes such as chromosomes 16, 25, 30, 31, 32, and 33, adding a total of 421 coding genes (**Supplementary Fig. 24f**, **Supplementary Note**).

**Figure 4.**
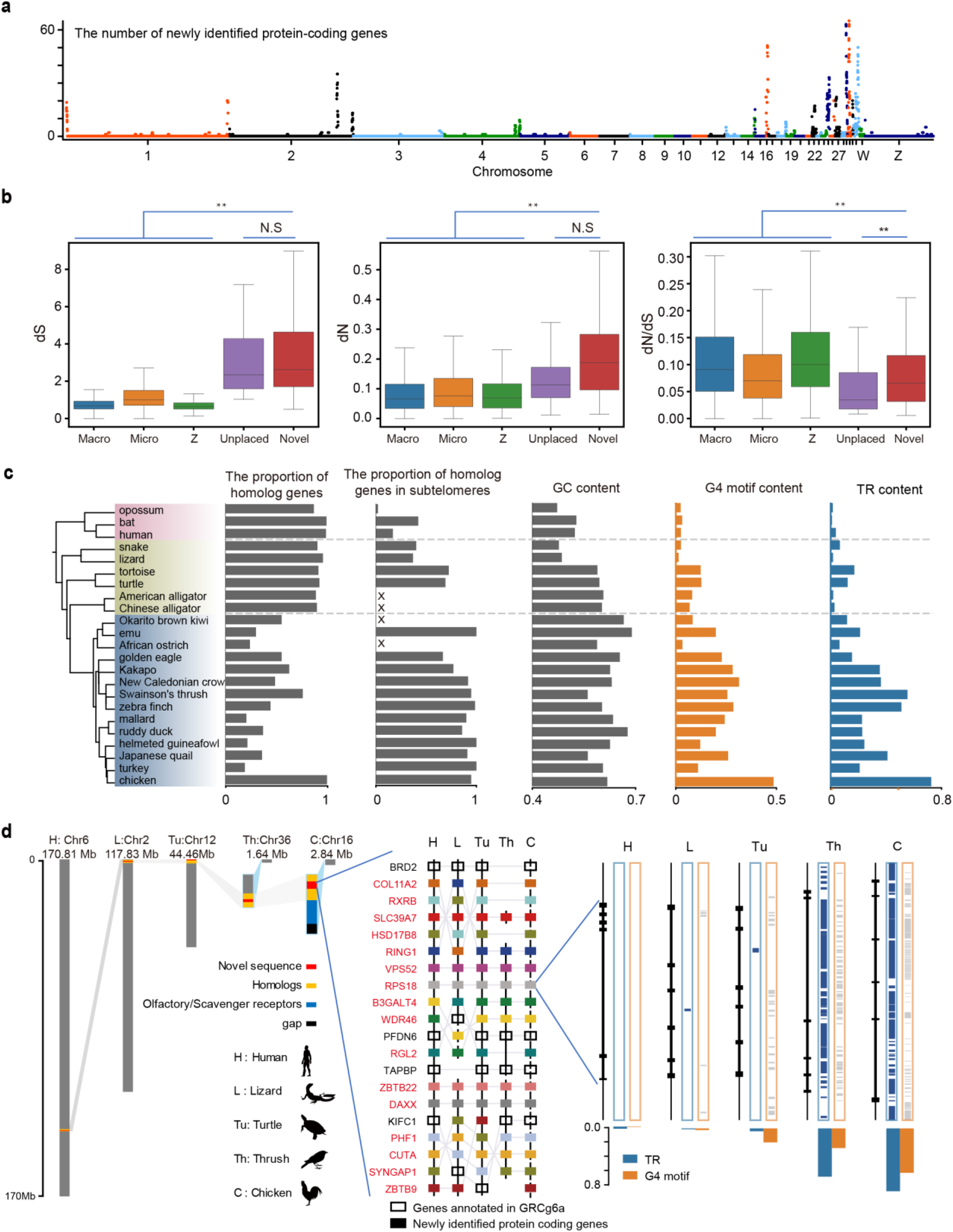
The novel coding genes clustered in the sub-telomere of chromosomes. **a**, The location of the novel coding gene clusters on chromosomes. **b**, Box plot for dS, dN, dN/dS values of genes on macro-chromosomes, micro-chromosomes, the Z chromosome, unplaced scaffold of chicken reference genome, and the novel sequences. **c**, The proportion of homologous gene number, homologous gene in sub-telomere, and contents of GC, G4 motif and TR of homologous novel coding genes clusters in chicken and 22 other species. The region containing more than 3 genes are considered as cluster, and genes located within 5-MB of the end of chromosomes are considered as sub-telomeric region. *d*, A detailed synteny conservation of novel coding genes on chromosome 16 of chicken with Mammal (human), Reptilia (lizard, turtle) and Aves (thrush, chicken), respectively. Hollow rectangles represent annotated genes in the genome, and other color rectangles, with gene name in red, represent novel coding genes in chickens.

It is widely accepted that sub-telomeric regions of the chromosomes and micro-chromosomes of birds exhibit high rates of recombination and mutation (International Chicken Genome Sequencing 2004; Burt 2005; Linardopoulou et al. 2005; Bell et al. 2020). We investigated the evolutionary rates of 160 high-quality orthologues of the novel coding genes by comparing chicken genes with those of human and mouse. The synonymous substitution rate (dS) and nonsynonymous substitution rate (dN) of these novel genes were 3.3- and 2.5-fold higher than that anchored GRCg6a genes, respectively. And the dN/dS ratio of these novel genes was lower than that of the reference genes. Interestingly, the unlocalized genes of GRCg6a, which may also be located in sub-telomeric regions, showed the similar mutation pattern as the novel genes (**Fig. 4b**). This suggested that the novel coding genes in sub-telomeric regions showed a higher mutation rate.

We next identified novel gene clusters to investigate collinearity. Screening according to the existence of more than 3 novel coding genes within 1 Mb bin across the genome revealed 19 regions containing 201 of 388 fully anchored genes. The 19 gene clusters were all located in sub-telomeric regions (**Fig. 4a**, **Supplementary Fig. 25**). Gene homology analysis indicated that almost all 201 novel coding genes had homologs in mammalian and reptile genomes and showed good collinearity (**Fig. 4d**, **Supplementary Fig. 25**, **Supplementary Table 13**). Some novel gene clusters likely existed in the sub-telomeric regions before the divergence of testudines and avian. However, the significant increase of the TR clusters with high content of noncanonical DNA structures only happened on the bird lineage (**Fig. 4c**). Unlike the previous notion that large segmental deletions occurred in the evolution process (Lovell et al. 2014; Zhang et al. 2014), our results provided a large number of confident new gene clusters in sub-telomeric regions, filling gaps in which genes were often missing due to insufficient sequencing.

### Functional assignment of novel regions and genes

Among the novel coding genes that we identified, 176 of them were identified as housekeeping genes in human and mouse (Hounkpe et al. 2021) (**Supplementary Table 9**). Through the annotation and enrichment analyses, we also found that a large number of them were involved in essential biological reactions and pathways, such as metabolism, signal transduction, basic biological functions, the immune system and disease (**Fig. 5a**, **Supplementary Tables 14**, **15 and 16**).

**Figure 5.**
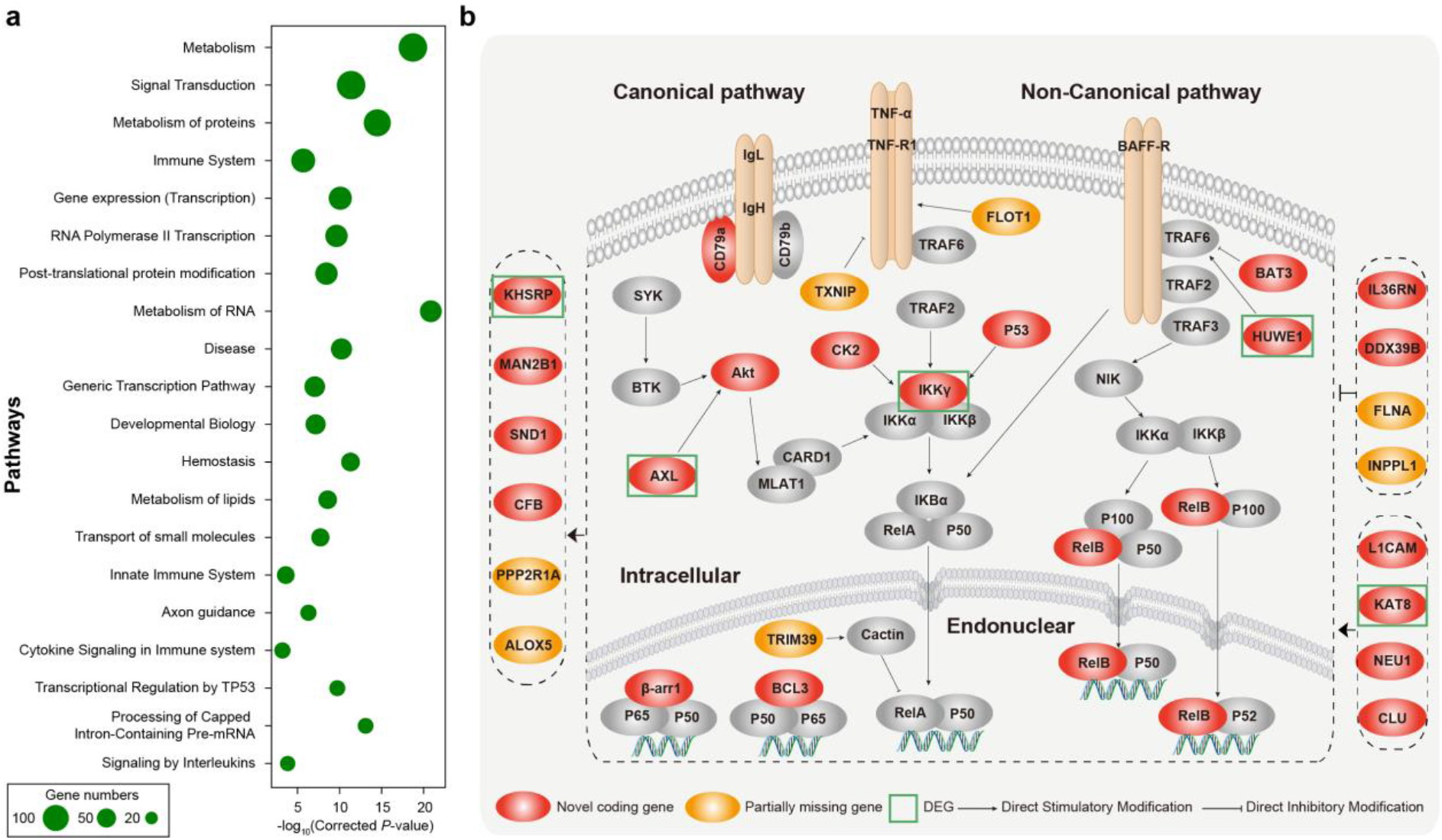
Function enrichment of novel coding genes and the case of NF-κB pathway related novel coding genes. **a**, The top 20 significant Reactome pathway with the largest number of novel coding genes. **b**, Novel coding genes (red) and partially missing gene (yellow) related to NF-κB signaling pathway. The green boxes represent differentially expressed genes (DEGs) in avian influenza virus.

In the novel regions, we dissected chromosome 16 and the sub-telomeric part of chromosome 1 as two examples to reveal their plausible gene arrangement and functions. Chromosome 16 is a micro-chromosome that contains many immune system-related genes (**Fig. 4d**) and spans only 2.84 Mb of GRCg6a. We assembled 3.76 Mb of novel sequences and identified 61 novel coding genes and 80 lncRNA genes on chromosome 16. The novel gene clusters showed good syntenic relationships with other tetrapods (**Fig. 4d**). One of the novel gene clusters showed that birds had experienced regional complications in the cluster and lacked a large number of coding genes (**Fig. 4d**). One novel coding gene, the complement factor B (*CFB*) gene, which is an important immune gene involved in the alternative complement pathway of the immune system (**Supplementary Fig. 27**) and is regulated by the nuclear factor kappa B (NF-κB) pathway, was *de novo* identified on chromosome 16. This gene is highly and uniquely expressed in the liver of chickens and confirmed based on our MS/MS data (**Supplementary Fig. 27**). In addition, we identified two novel ribosomal genes, mitochondrial ribosomal protein S18B (*MRPS18B*) and ribosomal protein S18 (*RPS18*) on chromosome 16 (**Fig. 4d**, **Supplementary Table 13**).

The other novel gene cluster, including the *leptin* gene, located on chromosome 1 (**Supplementary Fig. 25**). Based on RNA-Seq, previous research has shown that the *leptin* gene does exist in the chicken genome, yet it was absent from the chicken reference genome (Seroussi et al. 2017). Interestingly, we found that two divergent haplotypes of the *leptin* gene were assembled from two individuals. The entire gene region and its flanking regions had extremely high TR and G4 motif contents (**Supplementary Fig. 28**). Based on chromosome interaction data, *leptin* was assigned to the distal tip of chromosome 1p, showing collinearity with *SND1* and *LRRC4* (**Supplementary Fig. 25**, **Supplementary Fig. 28**). We found that *leptin* exon 2 was conserved, while exon 1 was variable in chicken. The length of its intron also varied among different chicken individuals (**Supplementary Fig. 28**). Neither of the two exons showed good homology with other species. In this region, we found another novel gene, ovocleidin-17 (*OC-17*), which plays a key role in avian eggshell biomineralization and is not contained in the reference genome (**Supplementary Table 9**).

### Application of the chicken pan-genome in avian influenza

The chicken pan-genome identify more crucial novel genes related to avian diseases resistance that had not been discovered previously. Chickens are susceptible to several diseases that have far-reaching effects on human society, such as avian influenza. Here, we reanalyzed the transcriptome data (Smith et al. 2015) of chicken lung and ileum samples after infection with low pathogenic (H5N2) and highly pathogenic (H5N1) avian influenza virus. Compared with the expression levels observed in the control group, 30 novel coding genes, 65 novel lncRNAs and 79 partially missing genes showed differential expression in these samples (FDR < 0.05) (**Supplementary Tables 9**, **10** and **12**). B cell-related genes (*CD22*, *CD79A*, *PRMT1*, and *SND1*), T cell-related genes (*CD2BP2*), immunoglobulin genes (*IGLL5*), and ribosome genes (*RPS18*) were screened among these differential expression genes (**Supplementary Tables 9** and **17**). Notably, several significantly differentially expressed genes (*AXL*, *HUWE1*, *IKKγ*, *KAT8*, and *KHSRP*) belonged to or were regulated by the NF-κB signaling pathway, which is the master regulator of the immune response to infection due to its role in regulating cytokine and antimicrobial peptide expression (**Fig. 5b**). *RELB*, a subunit of NF-κB, associated with the immune responses to influenza A (Ruckle et al. 2012) and severe acute respiratory syndrome-associated coronavirus (SARS-CoV) (Chen et al. 2006), was identified, anchored and validated in our study (**Supplementary Fig. 29**). Another novel gene, *IKKγ* (**Supplementary Table 9**), a subunit of the IκB kinase complex (IKK), was essential for the activation of NF-κB transcriptional activity. Besides, the newly identified genes *AXL* (Schmid et al. 2016), *CSNK2B* (Marjuki et al. 2008), *DDX39B* (Wisskirchen et al. 2011), *KHSRP* (Liu et al. 2015) and *TP53* (Wang et al. 2018) have also been reported to play a role in the immune response to influenza A. In total, there were 21 novel coding genes and 7 partially missing coding genes that belonged to or were regulated by the NF-κB signaling pathway (**Fig. 5b**). The NF-κB pathway is essential in defense against viral infections, such as those caused by influenza viruses.

## Discussion

The chicken is the modern descendant of the dinosaurs with its genome firstly sequenced in non-mammalian amniotes (International Chicken Genome Sequencing 2004). Despite several major updates, the completeness of the chicken genome still needs to be improved and the gene number of chicken is still argued. Here, our study suggests that the chicken pan-genome exhibits a more complex mammalian-like closed pattern. More specifically, we identified 1,335 and 3,011 novel coding genes and novel long noncoding genes, respectively, containing mostly core genes, which appear different from previous mammalian pan-genome studies that reported fewer novel genes (Sherman et al. 2018; Golicz et al. 2019; Sherman and Salzberg 2020; Tian et al. 2020). The highly complex noncanonical DNA structure across the novel genes might be the main reason to prevent the efficient genome assembly of identified novel genes in lots of individuals. The accidentally detection of DNA sequencing in the regions of novel sequences due to the secondary DNA structure might be the reason why there are still so many genes missing in the recent high-quality VGP avian assemblies, that may suggest there are still more challenges to complete the avian reference genome. Nevertheless, we increased the number of protein coding genes in chicken to 19,223 and denied the gene loss hypothesis during avian evolution. Although some genes may still hide in some more complex genomic regions and waiting to be discovered. Our study not only revealed the gene number in birds is comparable to that in other tetrapods but also presented a novel closed pattern of avian pan-genome. The complete avian genomes will greatly improve the studies on comparative genomics and functional genomics research in birds.

It has been believed that both the evolutionary substitution rate and the rate of chromosomal rearrangement in the avian lineage are lower than that of mammals (Burt et al. 1999; Zhang et al. 2014). However, we found that a large number of novel genes that have three times the substitution rate than the known ones, which can greatly increase the average substitution rate of the chicken genome. We find that the novel sequences and genes were concentrated in the sub-telomeric regions of chromosomes, in which the recombination rate tend to be higher (Linardopoulou et al. 2005; Bell et al. 2020). This may drive the base composition evolution via biased gene conversion (Marais 2003) and cause repeat expansions or contractions (Richard and Paques 2000; Polleys et al. 2017), and maybe the critical factor driving the development of the special characteristics in sub-telomeric regions. These genes may have a pivotal role on the formation and development of some unique phenotypes of the dinosaurs-avian branch. For instance, some differentially expressed novel genes were associated with immune response, which may be an ingenious design of the bird immune system to resist viruses with high mutation rates. With the high recombination rate, the novel sequences may represent a large unexplored party of the chicken genetic map, which will contribute to the comprehensive understanding of genetic variation and pinpoint the causal variations of important traits and thus promote the development of chicken breeding.

In conclusion, our chicken pan-genome provides a comprehensive resource and a great platform for the research of avian evolution, functional genomics, and chicken breeding. These results highlight the complexity of species genomes and suggest that many functionally important regions may be cryptic in reference genomes across the tree of life.

## Materials and Methods

### Sample collection

A total of 20 chicken individuals were collected from all around the world for genomic sequencing. Transcriptome sequencing was also performed in 11 tissues of six individuals, including breast muscle, bursa of Fabriclus, cecal tonsil, Harderian gland, hypophysis, hypothalamus, liver, ovary, spleen, testis, and thymus tissues. Moreover, tandem mass spectrometry (MS/MS) data were generated from three tissues (hypothalamus, spleen, and cecal tonsil) from four of the six individuals by RNA-Seq. The tissue sources and the institutes in charge of the collection are listed in **Supplementary Table 1.** All animal specimens were collected legally in accordance with the policies for the Animal Care and Use Ethics of each institution, making all efforts to minimize invasiveness.

### Library construction and genome sequencing

For PacBio continuous long reads (CLR) sequencing, genomic DNA was extracted from chicken liver using a QIAamp DNA Mini Kit (QIAGEN). The integrity of the DNA was determined with an Agilent 4200 Bioanalyzer (Agilent Technologies, Palo Alto, California). Eight micrograms of genomic DNA were sheared using g-Tubes (Covaris), and concentrated with AMPure PB magnetic beads. Each SMRT bell library was constructed using the Pacific Biosciences SMRTbell template prep kit 2.1. The constructed libraries were size-selected on a BluePippin™ system for molecules ≥ 20 kb, followed by primer annealing and the binding of SMRT bell templates to polymerases with the DNA/Polymerase Binding Kit. Sequencing was carried out using P6-C4 chemistry on the Pacific Bioscience Sequel II platform by Annoroad Gene Technology Company.

For short-read DNA sequencing, the genomic DNA of ten samples used for next-generation sequencing (NGS) assembly was extracted from ethylenediaminetetraacetic acid (EDTA)-anticoagulated blood randomly fragmented. Two paired-end libraries and two mate-pair libraries with insert sizes of 500 bp, 800 bp, 3 Kb and 5 Kb were constructed. All libraries were sequenced on the Illumina HiSeq 2000 platform according to the manufacturer’s protocol. After filtering out adapter sequences and low-quality reads, a total of 1.61 Tb (average 134 × coverage of chicken genome) of data were retained for assembly. In addition, the libraries of ten samples used for PacBio sequencing were also constructed using an amplification-free method with an insert size of 350 bp and sequenced on the Illumina XTen platform with paired-end 150 bp sequence reads.

### Whole-transcriptome sequencing

For transcriptome analysis, total RNA was extracted using TRIzol extraction reagent (Thermo Fisher). The RNA quality analysis method was the same as DNA quality analysis method described above. Libraries with 250-350 bp insert sizes were prepared using the TruSeq RNA Sample Prep Kit v2 (Illumina, San Diego, CA, USA). To obtain transcriptome profiles, all libraries were sequenced on Illumina XTen system platform using the manufacturer’s protocol.

### High-throughput chromatin conformation capture (Hi-C) sequencing

Hi-C experiments were performed according to a previously published protocol (Lieberman-Aiden et al. 2009). Hi-C libraries were created from the breast muscle samples of six of the above individuals. All libraries were sequenced on an Illumina HiSeq X Ten sequencer (paired-end sequencing with a 150 bp read length). On average, 127 Gb of data with approximately120-fold genomic coverage and 271,268,477 read pairs could be uniquely aligned to the chicken genome reference sequence (**Supplementary Table 2, Supplementary Table 3**).

### Tandem mass spectrometry analysis

The samples were ground into a cell powder in liquid nitrogen and then sonicated in lysis buffer (8 M urea, 1% protease inhibitor cocktail) three times on ice using a high-intensity ultrasonic processor (Scientz). The remaining debris was removed by centrifugation at 12,000 g at 4 °C for 10 min. Thereafter, the supernatant was collected, and the protein concentration was determined with a BCA kit according to the manufacturer’s instructions. Then, the protein solution was subjected to trypsin digestion. Next, the tryptic peptides were fractionated by high-pH reverse-phase HPLC using a Thermo Betasil C18 column (5 μm particles, 10 mm ID, and 250 mm length).

The tryptic peptides were dissolved in 0.1% formic acid (solvent A) and directly loaded onto a homemade reversed-phase analytical column (15-cm length, 75 μm i.d.). The gradient consisted of an increase from 6% to 23% solvent B (0.1% formic acid in 98% acetonitrile) over 26 min, an increase from 23% to 35% over 8 min and then to 80% over 3 min, with holding at 80% for the last 3 min, all at a constant flow rate of 400 nL/min in an EASY-nLC 1000 UPLC system. The peptides were introduced to a nanospray ionisation (NSI) source, followed by MS/MS in a Q ExactiveTM Plus system (Thermo) coupled online to the UPLC system.

The MS/MS data were processed using the MaxQuant search engine (v.1.5.2.8) (Cox and Mann 2008). Tandem mass spectra were searched against the human UniProt database concatenated with the reverse decoy database. Trypsin/P was specified as the cleavage enzyme, allowing up to 4 missing cleavages. The mass tolerance for precursor ions was set as 20 ppm in the first search and 5 ppm in the main search, and the mass tolerance for fragment ions was set as 0.02 Da. Carbamidomethyl on Cys was specified as a fixed modification and acetylation modifications and oxidation on Met were specified as variable modifications. The FDR was adjusted to < 1%, and the minimum score for modified peptides was set as > 40. For protein identification, peptides containing a minimum of seven amino acids and at least one unique peptide were required. Only proteins with at least two peptides and at least one unique peptide were considered to have been identified and used for further data analysis.

### *De novo* genome assembly, evaluation and repeat annotation

#### Assembly based on PacBio SMRT sequencing platform

The raw PacBio SMRT reads were corrected by itself with Canu v1.7 (Koren et al. 2017), and assembled with WTDBG v2.2 (Ruan and Li 2019) to generate the contig layout and edge sequences, and WTPOA-CNS v1.2 was used to obtain the initial consensus in FASTA format. Then, we used minimap2 v2.14-r883 (Li 2018) to map the corrected reads to the consensus, and they were subsequently polished by using WTPOA-CNS v1.2. This process was repeated three times. Next, the consensus sequence obtained in the previous step was mapped by using the NGS reads from the same individual with BWA-MEM v0.7.17-r1188 (Li and Durbin 2010) and then polished with Pilon v1.22 (Walker et al. 2014). This process was repeated three times to obtain the final contigs.

We performed further scaffolding based on the results for six individuals with Hi-C data. Using the final contigs as a reference, we mapped the Hi-C data to the final contigs using Juicer v1.5 (Durand et al. 2016) to obtain the interaction matrix. Finally, 3d-dna v180419 (Dudchenko et al. 2017) was used for scaffolding contigs.

#### Assembly based on NGS platform

The genomes sequenced on the NGS platform were *de novo* assembled into contigs by using a pipeline that combined the Fermi package (Li 2012) and Phusion assembler (Mullikin and Ning 2003) for 500 bp/800 bp paired-end libraries. For the 3 Kb/5 Kb mate-pair libraries, we used SOAPdenovo (Li et al. 2010) with 77 kmers to build contigs. Furthermore, SSPACE (Boetzer et al. 2011) was used to build scaffolds, and the contigs assembled by Fermi and Phusion were used for the substitution of sequences and bases and for further rectifying to rectify the local assembly error. After the inspection of the initial scaffolds, gaps were closed using Gap5 (Bonfield and Whitwham 2010) software.

#### Genome evaluation and annotation

BUSCO v3.0.2 (Simao et al. 2015) was used to assess assembly completeness by estimating the percentage of expected single-copy conserved orthologues captured in our assemblies and reference genome, referring to the lineage dataset aves_odb9 (Creation date: 2016-02-13, number of species: 40, number of BUSCOs: 4,915). Repeat sequences were annotated using RepeatMasker v4.0.8 (with the parameters: -engine ncbi -species ‘Gallus gallus’ -s -no_is - cutoff 255 -frag 20000). Subsequently, tandem repeats were further annotated using Tandem Repeats Finder v4.07b (Benson 1999) (with the settings 2 7 7 80 10 50 2000 -d -h). In addition, Quadron software (Sahakyan et al. 2017) was used to predict G4 motifs, and only nonoverlapping hits with a score greater than 19 were used for subsequent analysis.

### Chicken pan-genome construction

The *de novo* assemblies were aligned to the chicken reference genome (GRCg6a; GCF_000002315.6) using minimap2 (Li 2018) (-cx asm10). Based on the pairwise alignment, unaligned or low-identity sequences (showing more than 10% sequence divergence relative to GRCg6a) were extracted. Then, the adjacent sequences within 200 bp were merged. BLASTN 2.6.0+ (Camacho et al. 2009) (with the parameters -word_size 20 -max_hsps 1 - max_target_seqs 1 -dust no -soft_masking false -evalue 0.00001) was further used to align the unaligned sequences from the previous step to GRCg6a, and the sequences showing identity greater than 90% to GRCg6a sequences were removed. The remaining sequences were merged according to the adjacent regions within 200 bp, and sequences of less than 500 bp in length were removed. Subsequently, the unaligned and low-identity sequences obtained from all of the assemblies were combined, redundancies were removed with CD-HIT v4.7 (Fu et al. 2012) (parameter: -c 0.9 -aS 0.8 -d 0 -sf 1), and the longest sequence in the cluster was selected as the representative sequence. To further exclude potential contaminants in the dataset, we used BLASTN to compare the non-redundant set with the nr database of NCBI (v20181220). The sequences that were aligned to non-Chordata species were removed from the final novel sequence set (**Supplementary Table 5)**.

### Observed present or absent analysis of novel sequences in resequenced individuals

The whole-genome resequencing data of 922 chickens (Li et al. 2017; Wang et al. 2020) (**Supplementary Table 6**) were downloaded for the present or absent analysis of novel sequences. To explore whether the different sequencing platforms affected the results, the Illumina sequencing reads of the GRCg6a individual (SRR3954707 (Warren et al. 2017), which were previously used for single-base error correction) were also included in this analysis. The presence and absence of each novel sequence was then determined according to the sequence coverage and depth. First, to obtain high-quality reads and minimize false genotyping results due to low-quality reads supplied by Illumina, we implemented the following quality control procedures to filter the reads before read mapping using Trimmomatic v0.36 (Bolger et al. 2014), and leading or trailing stretches of Ns and bases with a quality score below 3 were trimmed. Then, the reads were scanned using a 4-base wide sliding window and clipped when the average quality per base was below 15, and only reads of 40 nucleotides or longer were finally retained. Second, high-quality paired reads were aligned to GRCg6a using BWA-MEM v0.7.17 (Li and Durbin 2010) with the default parameters, except that “-M” was enabled. The BWA-aligned BAM files were then processed using Picard v2.1 (http://broadinstitute.github.io/picard/), including reads sorted and merged read groups belonging to the same sample and marked duplicates at the sample level. Finally, we estimated the coverage distribution at each called site for each sample using QualiMap v2.2 (Okonechnikov et al. 2016).

Poorly aligned or unaligned reads were extracted as follows: Samblaster v0.1.24 (Faust and Hall 2014) was used to extract clipped reads and unaligned reads, while sambamba v0.6.8 (Tarasov et al. 2015) and SAMTools v1.9 (Li et al. 2009) were used to collect other poorly aligned reads. The paired reads with unaligned/poorly aligned read pairs were extracted using seqtk v1.3-r106 (https://github.com/lh3/seqtk) and were then aligned to the novel sequence set using a previously described process. Novel sequences with a coverage above 0.8 and a depth greater than one-quartered of the whole-genome depth were identified as present.

### Feature importance analysis

To estimate the influence of GC, G4 motif, and TR contents on the observed frequency of novel sequences, 9,200 novel sequences shared by all individuals were used to construct a random forest model. The sklearn package in Python was used to build the final model and perform classification.

### Transcribed region annotation and coding potential assessment

The raw RNA-Seq reads were processed to remove adapters, low-quality sequences and sequences with poly A/T tails using Trimmomatic v0.36 (Bolger et al. 2014). The cleaned reads were *de novo* assembled using SPAdes v3.14.1 (Bushmanova et al. 2019). The expression levels of the *de novo* assembled transcripts were quantified by using Kallisto v0.46.2 (Bray et al. 2016). Additionally, the cleaned reads were assembled using a reference-guided method by alignment to the *de novo* genome assemblies using HISAT2 v2.0.3-beta (Kim et al. 2019) with the default parameters, except that “--dta” was enabled. Transcripts including novel splice variants were assembled using StringTie v1.2.2 (Pertea et al. 2015) with the default parameters. Then, StringTie (--merge) was used to merge all the transcript GTFs obtained from the samples mapped to this assembly to obtain a reference annotation. Finally, all samples were reassembled and quantified using StringTie with the reference annotation to obtain the expression level of each transcript. Notably, the transcripts with fragments per kilobase per million mapped reads (FPKM) ≥ 1 were considered robustly expressed.

Redundancy among genes that were annotated based on the *de novo* and reference-guided methods and intersected with novel sequences was removed with CD-HIT (parameter: -c 0.9 -aS 0.8 -d 0 -sf 1). Then, the remaining genes were searched against the nr database and the genes of GRCg6a using BLASTN 2.6.0+. Genes with no hits to either non-Chordata species or GRCg6a were retained as “novel genes” that were completely absent in the chicken reference genome. Genes showing hits to GRCg6a genes with more than 95% identity were classified as partially missing in the chicken reference genome.

Next, the coding potential of these novel genes was assessed by using CPAT v1.2.3 (Wang et al. 2013) with the default parameters. CPAT uses an alignment-independent logistic regression model to detect coding potential based on sequence features. To select a cut-off for classification, we built hexamer tables and logit models for chicken using chicken CDSs and ncRNA sequences downloaded from Ensembl (release 98) as training data. Then, a two-graph receiver operating characteristic curve was used to determine the optimum cut-off value through 10 random sample validations (**Supplementary Figure 20**). A cut-off of 0.69 was selected to classify the novel genes as potential protein-coding or noncoding genes. Then, the ORFs were searched by using TransDecoder v5.5.0 (http://transdecoder.github.io) and ORFfinder v0.4.3 (https://www.ncbi.nlm.nih.gov/orffinder/) with the default parameters. Genes showing values above the cut-off of the CPAT with a minimum ORF of at least 100 amino acids were classified as novel coding genes. For the remaining novel genes, RNAcode (Washietl et al. 2011) was used to further estimate the coding potential. To prevent the divergent homologous haplotypes that can caused false gene duplications (Ko et al. 2021), we merged novel coding genes that have high similarity with each other or can be annotated to the same gene, and then performed manual check. We generated customized whole-genome alignments for each *de novo* assembly against Japanese quail (GCF_001577835.1), turkey (GCF_000146605.3), and helmeted guineafowl (GCF_002078875.1), which we used to estimate coding potential. We used BLASTX 2.6.0+ (with the parameters ‘-evalue 0.00001’) to translate each novel genes form all six possible reading frames, and the results were compared to known proteins in the nr database. Genes with an E value ≤ 10 ^-5^, alignment length of ≥ 10 amino acids and identity ≥ 95% were removed from the final potential long noncoding gene set. Only multiple exon genes with more than 200 nucleotides and without any detectable protein-coding potential were classified as novel long noncoding genes.

### Protein-coding gene annotation

Using the human (*Homo sapiens*) dataset as the background, the novel coding genes were annotated with the annotate module of online KOBAS 3.0 (Xie et al. 2011) (http://kobas.cbi.pku.edu.cn/). The Gene Ontology terms, KEGG pathways, and Reactome pathways of these genes were characterized by using the enrichment module of online KOBAS 3.0. *P* < 0.05 was set as the cut-off threshold.

InterProScan v5.36-75.0 (Jones et al. 2014) (parameter: -f tsv -dp) was used to classify the protein-coding gene fragments within the novel sequences and the protein-coding genes influenced by the location of novel sequences into protein families. The analysis results of Pfam 32.0 (http://pfam.xfam.org/) were selected to determine the families to which the proteins belonged.

### Differential expression analysis

The expression levels of each gene obtained from the previous step were used for differential expression analysis. The R language was used to identify differentially expressed genes with the edgeR package (v3.28.1) (Robinson et al. 2010). The fold changes between the two groups were calculated as logFC = log2 (experimental/control group). Benjamini-Hochberg correction was used to correct for multiple comparisons (with a false discovery cut-off of < 0.05). Genes in the two groups a with |logFC| > 2 and q-value < 0.05 were defined as differentially expressed genes.

### Anchoring novel sequences onto the reference genome

#### Flanking sequences

The novel sequences were anchored to GRCg6a based on alignment information between all *de novo* assemblies and GRCg6a. First, the scaffolds of the *de novo* assemblies that contained novel sequences were extracted and anchored on the chromosome/scaffold of GRCg6a which showed the most alignment hits with them. Then, the adjacent flanking sequences (more than 100 bp) of the novel sequences aligned to the same chromosome/scaffold were retained for further positioning. If the flanking sequences were perfectly aligned to GRCg6a with no gaps, an identity ≥ 90% and a breakpoint shift of ≤5bp, we recorded the sequences as “placed”. The other alignments were recorded as “ambiguously placed”. The novel sequences with two placed flanking sequences were reported as “localized”. The novel sequences with one or two ambiguously placed flanking sequences were reported as “unlocalized”. The final remaining sequences were reported as “unplaced”. Based on the genome placement information, the localized sequences could be further classified as insertions, alternate alleles, or ambiguous sequences. The insertions introduced only one sequence fragment to the reference genome and were no more than 10 bp in length. For alternate alleles, the novel sequences had to share less than 90% (or 0%) identity with their counterparts in the reference. Furthermore, the novel sequences and their counterparts had to have comparable lengths, with a length ratio between 1/3 and 3. The remaining sequences that did not meet the above criteria for insertions and alternate alleles were classified as ambiguous sequences.

#### Chromosome interaction mapping

The preprocessing of paired-end sequencing data, mapping of reads and filtering of mapped di-tags were performed using the Juicer pipeline (version 1.5) (Durand et al. 2016). Briefly, short reads were mapped to the chicken pan-genome using BWA-MEM (version 0.7.17-r1188) (Li and Durbin 2010). Reads with low mapping quality were filtered using Juicer with the default parameters, discarding invalid self-ligated and un-ligated fragments as well as PCR artefacts. Filtered di-tags were further processed with Juicer command line tools to bin ditags (10 kb bins) and to normalize matrices with KR normalization (Knight and Ruiz 2012). We normalized all Hi-C matrices on the same scale by KR normalization, ensuring that any differences between Hi-C data were not attributable to variation in sequence length. The maximum 100-kb bin of each novel sequence interaction (interaction intensity ≥5) was collected as a potential location of novel sequences. Novel sequences that were validated in at least two individuals with Hi-C data and anchored to the same location were remained.

### Gene orthology and dN/dS analysis

The integrated toolkit TBtools v1.0 (Chen et al. 2020) was used for collinearity analysis between species. First, the protein sequence of each gene was obtained, and pairwise sequence similarities were calculated using BLASTP with a cut-off of E value ≤ 10^-10^. Then, syntenic blocks were detected using MCScanX v1.0 (Wang et al. 2012) with the default parameters. OrthoFinder v2.4.0 (Emms and Kelly 2019) was used to identify orthologous genes with the default parameters. Among these genes, 1:1 orthologous genes between different species were used for downstream analysis. Using 1:1 orthologous genes as the input, Codeml in PAML version 4.9d (Yang 2007) was used for dN/dS analysis with the default parameters. The genome assemblies and corresponding annotations used in this analysis as below: Gray short-tailed opossum (GCF_000002295.2), Greater horseshoe bat (GCF_004115265.1), Human (GCF_000001405.39), Western terrestrial garter snake (GCF_009769535.1), Common lizard (GCF_011800845.1), Red-eared slider turtle (GCF_013100865.1), Goodes thornscrub tortoise (GCF_007399415.2), Green sea turtle (GCF_015237465.1), American alligator (GCF_000281125.3), Chinese alligator (GCF_000455745.1), Australian saltwater crocodile (GCF_001723895.1), Okarito brown kiwi (GCF_003343035.1), African ostrich (GCF_000698965.1), Emu (GCA_016128335.1), Golden eagle (GCF_900496995.1), Kakapo (GCF_004027225.2), New Caledonian crow (GCF_009650955.1), Swainson’s thrush (GCF_009819885.1), Zebra finch (GCF_008822105.2), Mallard (GCF_015476345.1), Helmeted guineafowl (GCF_002078875.1), Turkey (GCF_000146605.3), Japanese quail (GCF_001577835.2), and Chicken (GCF_000002315.6).

## Supplementary Materials

Supplementary Note

Supplementary Figure 1 to 29

Legends for Supplementary

Table 1 to 17

Supplementary References

## Acknowledgements

We thank High-Performance Computing (HPC) of Northwest A&F University (NWAFU) for providing computing resources. We thank Margarida Cardoso-Moreira for kindly providing the individual source information of the 217 chicken transcriptomes.

## Funding

This project was supported by grant from the National Natural Science Foundation of China (31822052, 31572381) and the National Thousand Youth Talents Plan to Y.J., National Natural Science Foundation of China (No. 31930105) and China Agriculture Research System of MOF and MARA(CARS-40) to N.Y., National Natural Science Foundation of China (31761143014) to Q.N. and the Unit of Excellence 2021, University of Phayao, Thailand (UoE64003) to C.S.

## Author contributions

N.Y., Y.J. and X.H. conceived the project and designed the research. M.L., X.T., P.B. and N.X. performed the majority of the analysis with contributions from Y.W., X.D., R.L., Y.G., F.W., Xiangnan W., P.Y., S.Z.. C. Sun, C.W., F.L., X.L., A.S. and C.S. prepared the DNA samples. X.J. and Q.N. provided the genome of Fayoumi. M.L., Xihong W. and N.X. drafted the manuscripts with input from all authors, and Y.J., C. Sun, Y.W., R.H. M.W. and X.Z. revised the manuscript.

## Competing interests

The authors declare no competing interests.

## Data availability

All the data of our study are publicly available at the NCBI Sequence Read Archive (https://www.ncbi.nlm.nih.gov/sra) under accession code BioProject: PRJNA573584. The data are available at https://dataview.ncbi.nlm.nih.gov/object/PRJNA573584?reviewer=buapp4f6dl06ls0snslqmitqh6 and http://animal.nwsuaf.edu.cn/code/index.php/panChicken/loadByGet?address[]=panChicken/Download/Denovo.php.

## Notes

https://dataview.ncbi.nlm.nih.gov/object/PRJNA573584?reviewer=buapp4f6dl06ls0snslqmitqh6

http://animal.nwsuaf.edu.cn/code/index.php/panChicken/loadByGet?address[]=panChicken/Download/Denovo.php

### Competing Interest Statement

The authors have declared no competing interest.

